# Variation in transformation frequency and competence gene expression among serotype 3 Streptococcus pneumoniae

**DOI:** 10.1101/2024.08.19.608640

**Authors:** Smitha Shambhu, Eleonora Cella, Taj Azarian

## Abstract

**Background:** Streptococcus pneumoniae is a human commensal and the causative agent of pneumococcal disease. Pneumococci are naturally competent – able to uptake exogenous DNA from the environment and incorporate it into their genome through homologous and non-homologous recombination. Recombination has significantly shaped the evolutionary history of S. pneumoniae, as it allows pneumococci to rapidly adapt. Recombination frequencies vary considerably across pneumococcal populations, yet underlying mechanisms for these variations are not well understood. Entry and exit into competence, a state in which the cell can uptake DNA, is tightly regulated through transcriptional changes.

**Methods:** To elucidate differences in transformation frequency among strains as well as the underlying genetic mechanisms, we carried out in-vitro competence assays and measured gene expression changes during the competent state using RNA sequencing of representative strains belonging to Serotype 3 clonal complex (CC) 180 and a non-CC180 comparison.

**Results:** We observed consistent differences in transformation frequencies among groups, which correlated with variation in differentially expressed genes during competence. While all strains exhibited a similar response to competence stimulating peptide (CSP) for early competence genes, we observed variation in expression of late competence genes, which encode the DNA uptake apparatus, DNA repair and recombination proteins. We also observed differences in expression of genes linked to bacteriocin production, which may partially explain observed population-level differences. Further genomic analysis suggests variation in promoter sequences governing late competence genes may be slowing transition from early to late components of the competence pathway.

**Conclusions:** We show that there is considerable variation in competence even among closely related strains belonging to the same CC and that this variation may be the result of subtle genomic differences. Additional studies are needed to assess the phenotypic impact of genomic variations.

## Introduction

Streptococcus pneumoniae is a human commensal and the causative agent of invasive pneumococcal disease (IPD). The capsular polysaccharide covering pneumococci determines the serotype and is the main virulence factor as well as the target of the pneumococcal conjugate vaccines (PCV) ^1^. Among the 100+ recognized serotypes, serotype 3, known for its mucoid capsular phenotype, is associated with higher mortality rates compared to other pneumococci ^2–4^. Despite its inclusion in the 13-valent pneumococcal conjugate vaccine (PCV13), serotype 3 remains a significant cause of IPD, and recent studies have also shown the emergence of serotype 3 as the most prevalent serotype in IPD after PCV-13 introduction, replacing serotype 19F ^5–7^.

While strains belonging to Global Pneumococcal Sequence Cluster (GSPC) 12 or clonal complex (CC) 180, as defined by multi-locus sequence type (MLST), are the most dominant serotype 3 lineage globally, a recent shift in the population structure within GPSC12/CC180 has been identified ^8–10^. Using whole-genome sequencing data, CC180 was resolved into two distinct phylogenetic clades, Clade I with subclades Clade I- α and Clade I- β, and Clade II. The persistence of serotype 3 as a significant cause of IPD corresponded with a global increase in the proportion of strains belonging to Clade II. Further, as compared to Clade I, Clade II strains were found to have a higher prevalence of antibiotic resistance elements and diverse non-capsular surface protein antigens, which was possibly driven by a higher rate of recombination as estimated through analysis of population genomic data ^10^.

Competence, or the ability to undergo homologous recombination, is one means by which S. pneumoniae can evolve in a relatively short period of time ^11^. For example, rapid pneumococcal evolution in response to vaccine induced selective pressure was observed after the introduction of PCVs, with the rise of non-vaccine serotypes as well as the identification of several ‘capsule switching’ events, whereby the entire cps locus is transferred to a different genomic background ^12–14^. Competence is also closely linked to pneumococcal virulence ^15^. In the competent state, pneumococci initiate cellular processes dedicated to acquisition of DNA from the extracellular environment, which include the release of allolysis toxins that incidentally increase virulence ^16,17^. Pneumococci vary in their ability to undergo homologous recombination as measured by population genomic approaches and through in-vitro studies ^18^. The latter induces competence using synthetic competence stimulating peptide (CSP) to measure the uptake of a selectable marker gene transferred to a recipient ^19–21^.

Competence is initiated by the binding of CSP, a quorum-sensing pheromone protein encoded by the gene comC, to its receptor encoded by comD which is a histidine kinase activating ComE through phosphorylation; initiating transcription at promoters with the ComE-box, a 9bp repeat element in the S. pneumoniae genome (Figure 1) ^22,23^. Genes induced by comE, known as the early com genes, include the comCDE, comAB operons as well as comX which encodes an alternative sigma factor ^24^. ComX induces transcription of genes encoding DNA processing proteins such as the single-stranded DNA binding protein SsbB and recombinase RecA ^25,26^ as well as genes important for virulence such as lytA, cbpD and comM ^27^. Exit from the competence state is thought to involve dprA, comW and comE which inhibit comCDE transcription by outcompeting ComE-P for binding to the comCDE promoter ^28,29^. The induction and exit from competence are under tight temporal regulation as revealed by studies measuring gene expression changes during competence ^30–32^(Figure 1).

**Figure 1.**
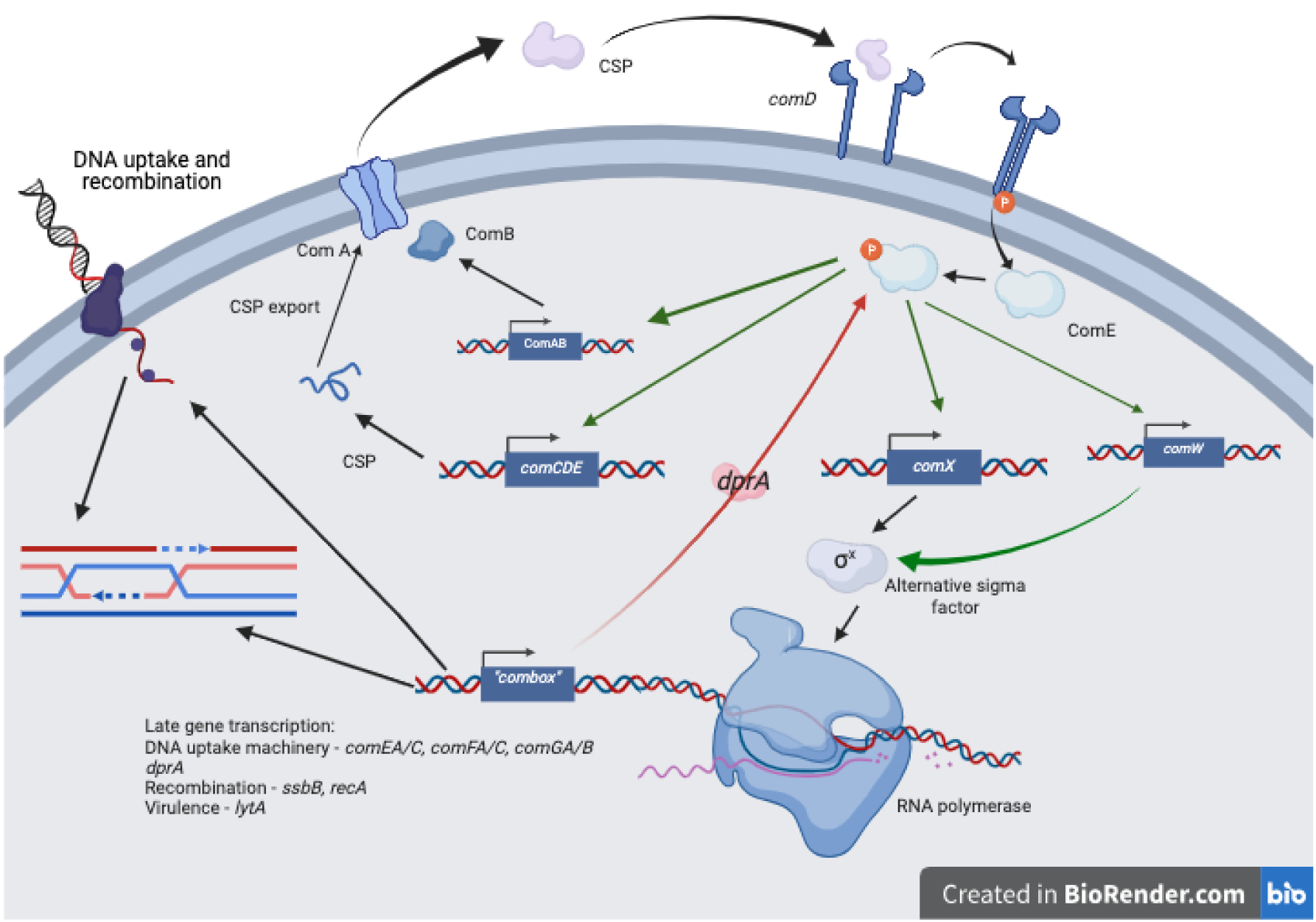
Diagram of competence pathway in Streptococcus pneumoniae.

Variation in competence within CC180 may explain some of the observed changes in the population genomics of S. pneumoniae serotype 3. Currently, little is known about the transformation frequencies of S. pneumoniae serotype 3 strains and the underlying genetic mechanisms responsible for differences in DNA uptake and recombination. In this present study, we sought to characterize phenotypic differences in competence between the two dominant clades of S. pneumoniae serotype 3 by measuring in-vitro transformation frequencies. Further, using genomic analyses and measuring differences in expression of competence genes, we aimed to identify factors associated with differences in competence.

## Methods

### Strains and Culture Conditions

To investigate phenotypic variation in competence between S. pneumoniae serotype 3 CC180 clades, we studied previously published strains representative of Clade I (n = 3) and Clade II (n = 3) compared to S. pneumoniae serotype 2 laboratory strain D39 ^10,33^. S. pneumoniae from -80℃ stocks were thawed and streaked onto trypticase soy agar plates with 5% (vol/vol) sheep blood, henceforth referred to as blood agar (BA) plates and grown overnight at 37℃ with 5% carbon dioxide. For inoculations, single isolated colonies were inoculated into Todd-Hewitt broth supplemented with 5g/L yeast extract (THY) and grown for 6h at 37℃ with 5% carbon dioxide.

### In-vitro transformation

S. pneumoniae from frozen stocks were thawed and streaked onto BA plates and grown in THY broth to an optical density of 0.4 at 620 nm (OD_620_), diluted 100-fold with fresh medium and grown to an OD_620_ of 0.05 in THY containing 0.5% (wt/vol) glycine (pH 6.8). 200 µl of the culture was pelleted by centrifugation at 12,000 rpm for 5 min at room temperature. The pellet was gently resuspended in 200 ul of THY containing 0.2% (wt/vol) bovine serum albumin (BSA), 0.5% nM CaCl_2_ and 0.5% (wt/vol) glycine (pH 8.3). The cultures were treated with 500 ng/ml of synthetic CSP-1 and incubated at 37℃ for 15 min. The cultures were then treated with an amplicon of the aphIII gene in the sweet janus cassette integrated into TIGR4 genomic DNA conferring resistance to Kanamycin at a concentration of 1 µg/ml ^34^. The aphIII gene was amplified with the primer pair TIGR4_KSJ_1. The mixture was incubated at 37℃ for 1h, mixed with 800 µl of THY and incubated for 2 additional hours before being serially diluted and plated on BA plates with 200 µg/ml of kanamycin. The plates were incubated overnight at 37℃ and colonies were counted to calculate the number of transformants. 100 µl of the same culture was serially diluted and plated onto BA to determine the total number of bacteria in the sample. Transformation frequency (TF) was calculated as follows. TF = (CFUK x DF) / (CFU x DF), where CFUK is the number of colony-forming units on the kanamycin plate, CFU is the number of colony-forming units on the antibiotic-free BA plate and DF is the dilution factor. Four replicates for three strains belonging to Clade I, three strains belonging to Clade II, and D39 were performed, and the log-transformed mean transformation frequency was calculated. Differences in transformation frequency among each group were analyzed by statistical analysis of variance (ANOVA) using R.

### RNA Extraction and Sequencing

To better understand the genotypic basis for variation in transformation frequencies, we further investigated global regulation of competence-associated genes through RNAseq transcriptomic analysis. We selected one strain from Clade I (PT8465) and Clade II (ND6401) as well as D39 based on transformation frequency values from the in-vitro transformation assay. Isolates from frozen stocks were thawed and streaked onto BA plates and grown overnight at 371 with 5% carbon dioxide. Individual colonies were then grown in THY broth to OD_620_ of 0.4.

Cultures were treated with 200 ng/ml of CSP. Two volumes of RNAprotect bacteria reagent (Qiagen) were added to the cultures at 10, 15, 20 mins after CSP addition, vortexed and incubated at room temperature for 5 mins. These timepoints were selected as it has been shown that peak competence is achieved after 15 - 20 mins ^30^. To determine baseline expression levels, a sample was taken at time 0 minutes just prior to CSP addition. Cultures were then centrifuged at 5000g for 10 mins. The pellet was resuspended in Buffer RLT and transferred to a MP biomedicals Lysing matrix B tube. Bacterial cells were disrupted in the TissueLyser (Qiagen) for 5 mins at 30Hz and supernatant was used as lysate input into the Qiagen RNeasy mini kit. Purified RNA was measured using Qubit and stored at -801. cDNA libraries were made using Illumina stranded RNA library preparation kits with RiboZero Plus rRNA depletion. Libraries were sequenced on NextSeq 550 to a target depth of 12M paired end reads (2x50bp) per sample. Each of these biological strains was run in three technical replicates for each timepoint.

### Analysis of RNA Sequencing data

Quality of RNA sequencing data was assessed using FastQC 0.11.9 and reads with low quality or high N content were filtered out using fastp with default settings. Filtered reads were then mapped to the S. pneumoniae serotype 3 SPN-OXC141 reference genome (NCBI GenBank: FQ312027.1) using Bowtie2 v2.4.5, and the number of reads mapped to each gene were counted using featureCounts. Minimal pre-filtering was performed according to recommendations from the DESeq2 manual wherein only genes that have at least 10 reads total across all samples were kept in the read counts tables. Read count files were imported into R and DESeq2 v3.16 was used to identify differentially expressed genes. The function plotPCA was used to generate PCA plots comparing the transcriptomes of each strain at different timepoints during competence. To compare trends during the stages of competence among strains, we plotted heatmaps of the top 100 differentially expressed genes using Pearson’s correlation coefficient and hierarchical clustering functions. We assessed the most highly upregulated genes during the competence process using volcano plots generated using ggplot2 with R comparing gene expression at 15 minutes post CSP addition to 0 mins. We classified competence genes as “early” or “late” by their role in the competence pathway ^30^ and assessed how expression of early and late competence genes varied at 15 minutes post CSP addition to 0 minutes by plotting their fold change and normalised counts. The RNA-Seq accession number is GEO Submission (GSE273581).

## Results

### In-vitro transformation frequency

We studied 6 strains of S. pneumoniae serotype 3 belonging to two distinct clades of CC180 and a non-serotype 3 comparison strain, D39. We found all strains to be competent as confirmed by their ability to uptake the kanamycin resistance marker. Serotype 3 isolates exhibited a higher transformation frequency than the serotype 2 strain D39. Within serotype 3, Clade I isolates demonstrated a consistently higher transformation frequency in comparison to strains belonging to Clade II. Differences in transformation frequency were statistically significant (p < 0.05) and varied by up to 2 orders of magnitude from 2.57 × 10^-5^ to 1.15 × 10^-3^ (Table 1 and Figure 2).

**Figure 2.**
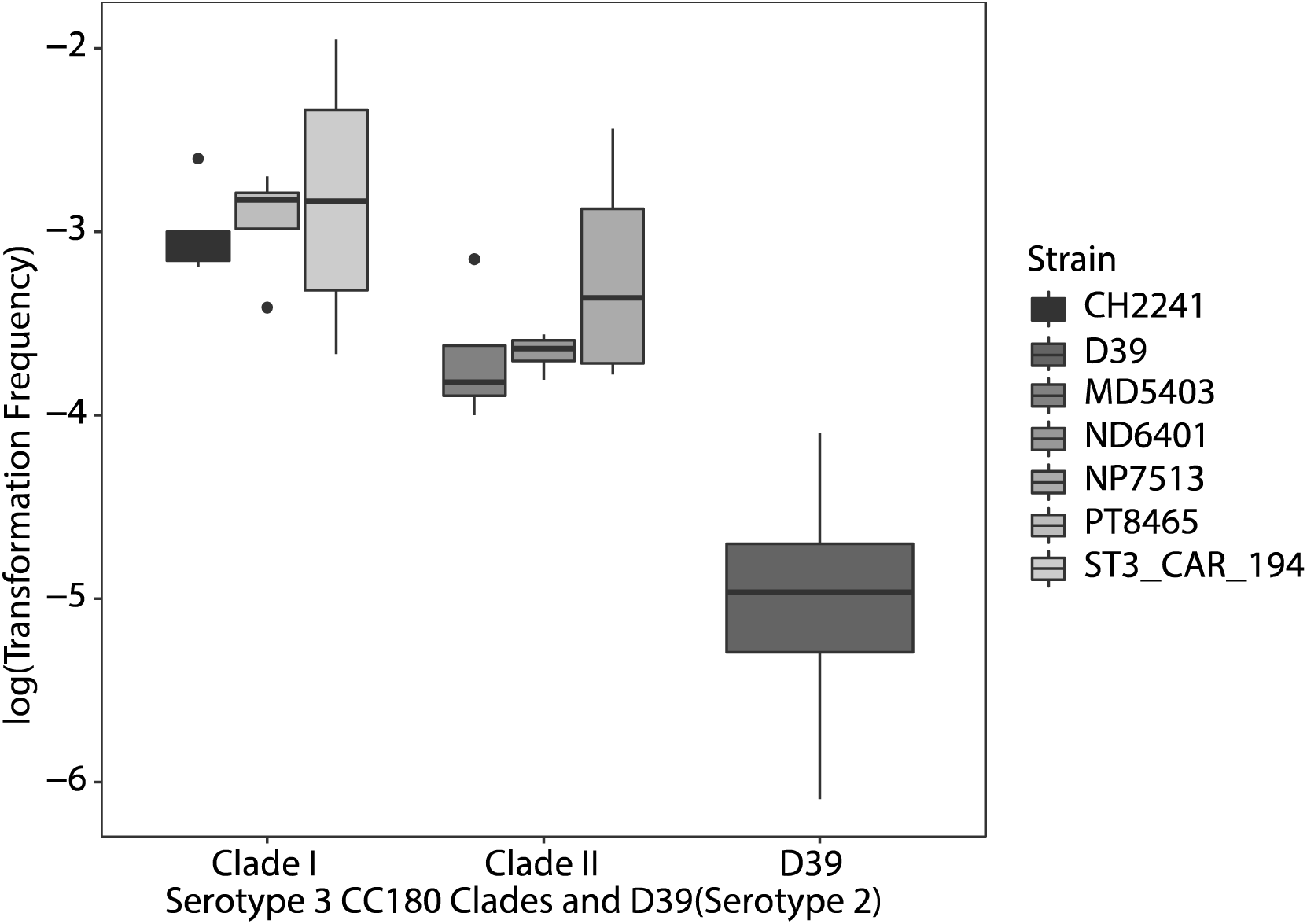
Transformation frequencies of S. pneumoniae serotype 3 isolates and D39. Transformation frequency is represented as log10 transformed values on the y-axis. Data plotted by taking mean of four replicates for each strain. Clade refers to population structure of serotype 3 CC180.

**Table 1.** Mean transformation frequency of serotype 3 strains and D39 calculated as number of transformants (CFU on kanamycin plate) / total number of colonies (CFU on blood agar). Mean calculated from 4 replicates.

| Strain | Clade | Mean Transformation Frequency | Mean ± S.D.<br>log(Transformation Frequency) |
| --- | --- | --- | --- |
| PT8465 | Clade I | 1.34E-03 | $-2.942 \pm 0.320$ |
| CH2241 | Clade I | 1.15E-03 | $-3.018 \pm 0.279$ |
| ST3_CAR_194 | Clade I | 3.85E-03 | $-2.822 \pm 0.761$ |
| NP7513 | Clade II | 1.24E-03 | $-3.234 \pm 0.630$ |
| ND6401 | Clade II | 2.24E-04 | $-3.660 \pm 0.109$ |
| MD5403 | Clade II | 2.78E-04 | $-3.697 \pm 0.376$ |
| D39 | NA | 2.57E-05 | $-5.030 \pm 0.819$ |

### Competence transcriptomes in S. pneumoniae

We obtained an average of 11 million (M) reads per sample (range: 8.1M - 13.5M). Upon filtering for read quality, we retained an average of 10.7M reads per sample. First, we used principal component analysis (PCA) to describe the general behavior of each timepoint for our pneumococcal strains. The analysis showed that replicates of each timepoint tend to cluster together (Supplementary Figure 1A-C). Timepoints after CSP addition when cells are in the competent state also tend to cluster together and away from the 0 min timepoint where cells have not been stimulated to enter competence. This indicated overlapping expression patterns of competence related genes across replicates.

**Supplementary Figure 1.**
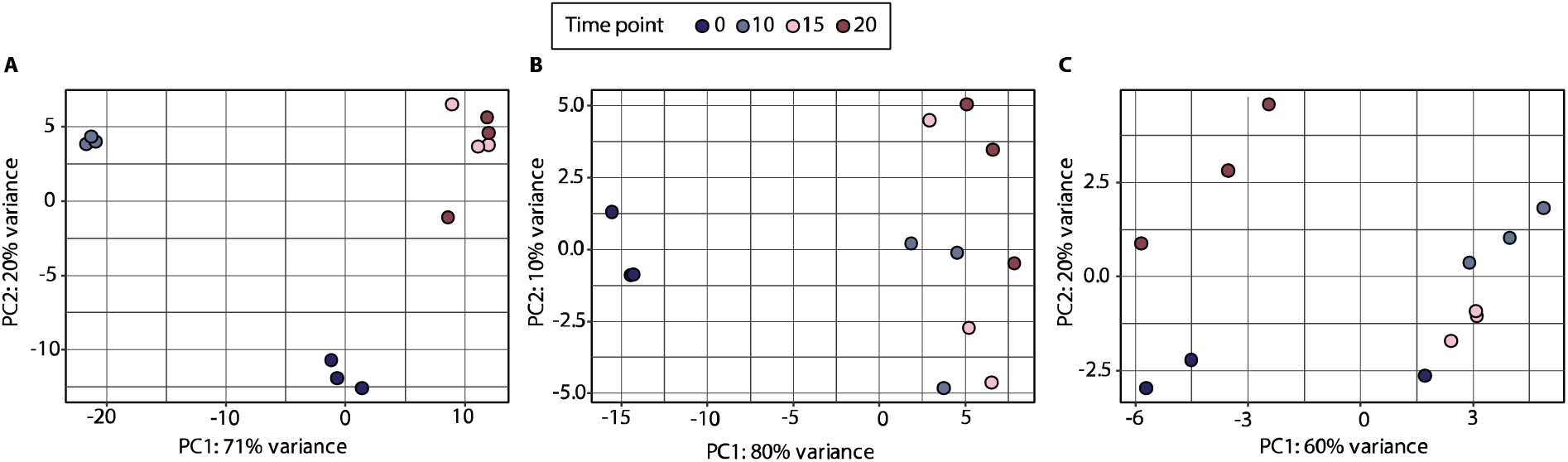
Principal component analysis of the normalized transcriptome profile of bacterial cells at different timepoints after CSP addition for D39 (panel A), PT8465 (Clade I) (panel B) and ND6401 (Clade II) (panel C). The different timepoints are colored according to the legend is on the top of the figure.

### Differential expression of competence-associated genes

To identify competence genes that were differentially expressed, we first screened our dataset for all genes that were significantly upregulated (p-value < 0.05, log_2_ (fold change) > 1.0) at 10, 15 or 20 minutes after CSP addition in comparison to the 0 minute timepoint. We focused on upregulated genes known to be associated with the competence cascade including comX and comE. D39 exhibited a larger number of upregulated genes (n = 341) in comparison to our Clade I and II serotype 3 strains (Table 2).

**Table 2.** Number of significantly upregulated (p-value < 0.05, log2(fold change > 1.0) early and late competence genes for each strain at the different timepoints measured.

| Timepoint | Strain |  |  |
| --- | --- | --- | --- |
|  | D39 | PT8465 (Clade I) | ND6401 (Clade II) |
| 10 mins | 341 | 72 | 13 |
| 15 mins | 122 | 85 | 9 |
| 20 mins | 131 | 85 | 10 |

To assess dynamic changes in expression of competence-associated genes over time, we plotted heatmaps of all known competence genes using regularized log (rlog) transformed counts from DESeq2 (Supplementary Figures 2-4). Among our Clade I (PT8465) and Clade II (ND6401) serotype 3 strains we observed a consistent upregulation of early competence genes starting at 10 mins after CSP addition including the comAB, comCDE operons and comX. While PT8465 shows consistent upregulation of the late competence genes such as the recA operon, comEA, comFA and comG operons, ND6401 does not exhibit the same expression pattern. D39 however showed upregulation of early and late competence genes across replicates at 15- and 20 minutes after addition of CSP. The blpY operon encoding bacteriocins show higher expression levels at 15 and 20 minutes for D39 and PT8465 but only at 20 minutes for ND6401. ND6401 does upregulate another bacteriocin peptide encoded by briC at 10, 15 and 20 minutes after CSP addition.

**Supplementary Figure 2.**
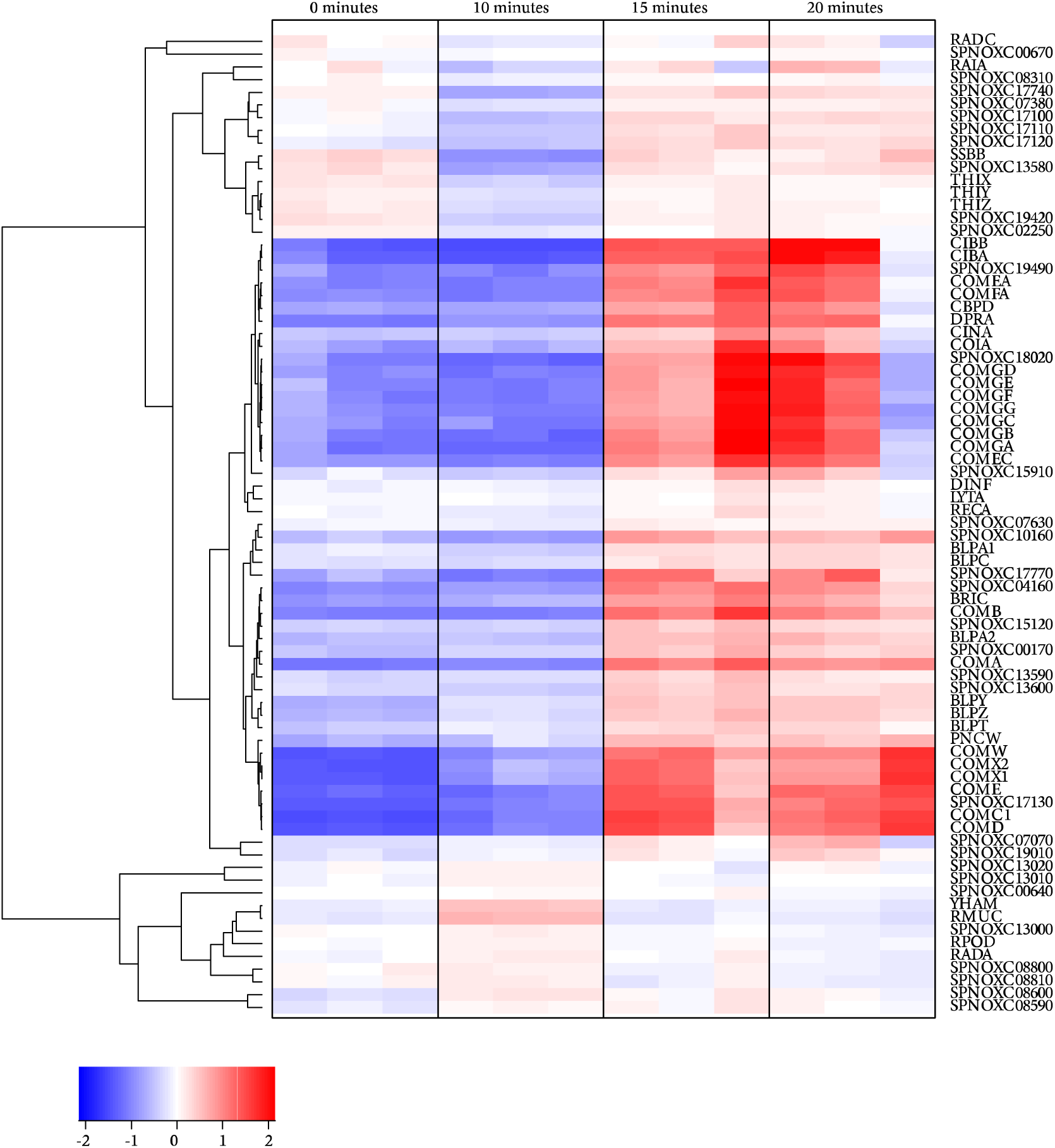
Heatmap showing levels of expression of known competence genes in D39 at 10, 15, 20 minutes after CSP addition. Samples are ordered by timepoint and replicate number on the x-axis while genes on y-axis are colored according to their z score and clustered with genes having a similar expression profile using hierarchical clustering. There are three replicates for each timepoint.

**Supplementary Figure 3.**
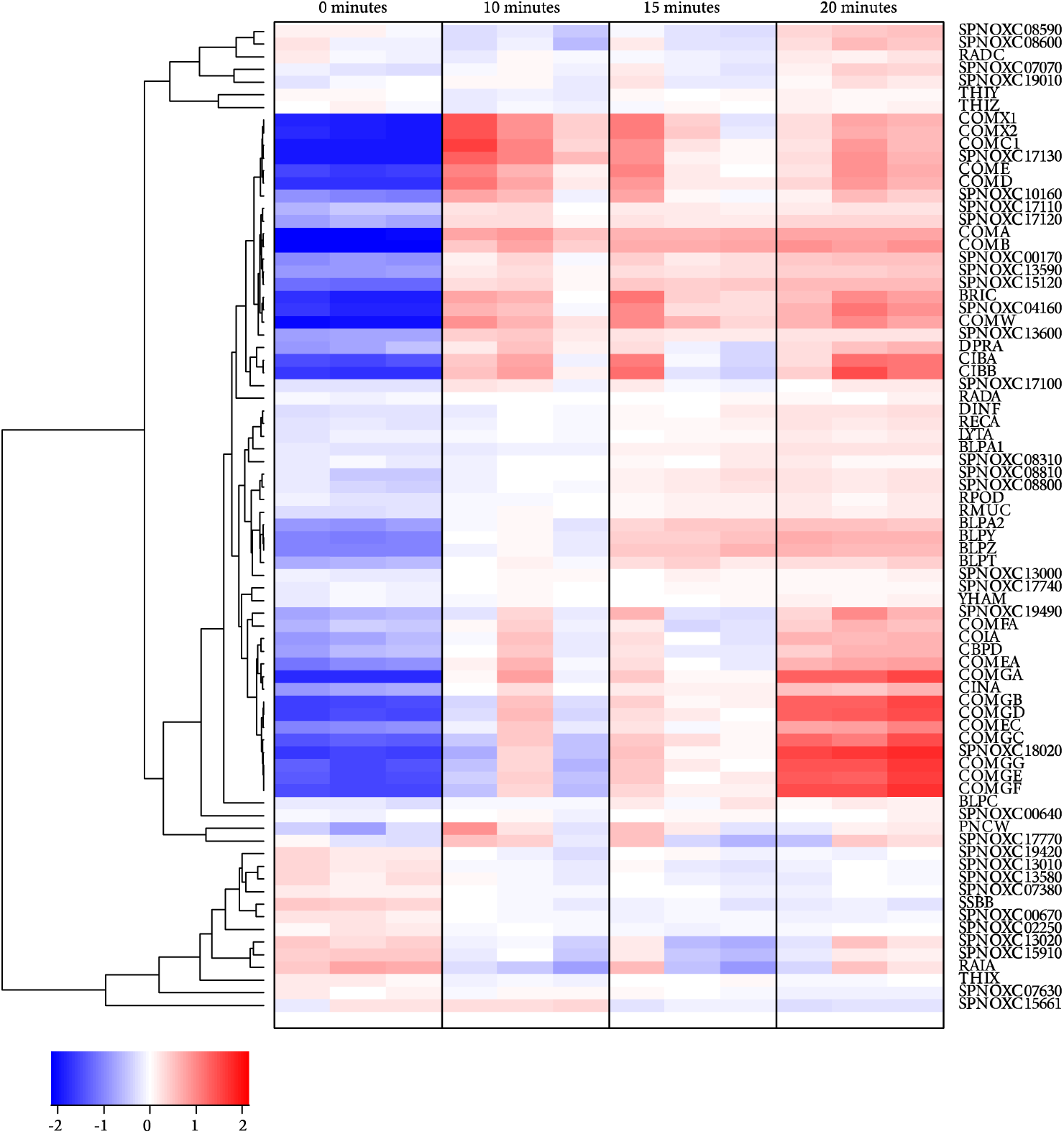
Heatmap showing levels of expression of known competence genes in PT8465 (Clade I) at 10, 15, 20 minutes after CSP addition. Samples are ordered by timepoint and replicate number on the x-axis while genes on y-axis are colored according to their z score and clustered with genes having a similar expression profile using hierarchical clustering. There are three replicates for each timepoint.

**Supplementary Figure 4.**
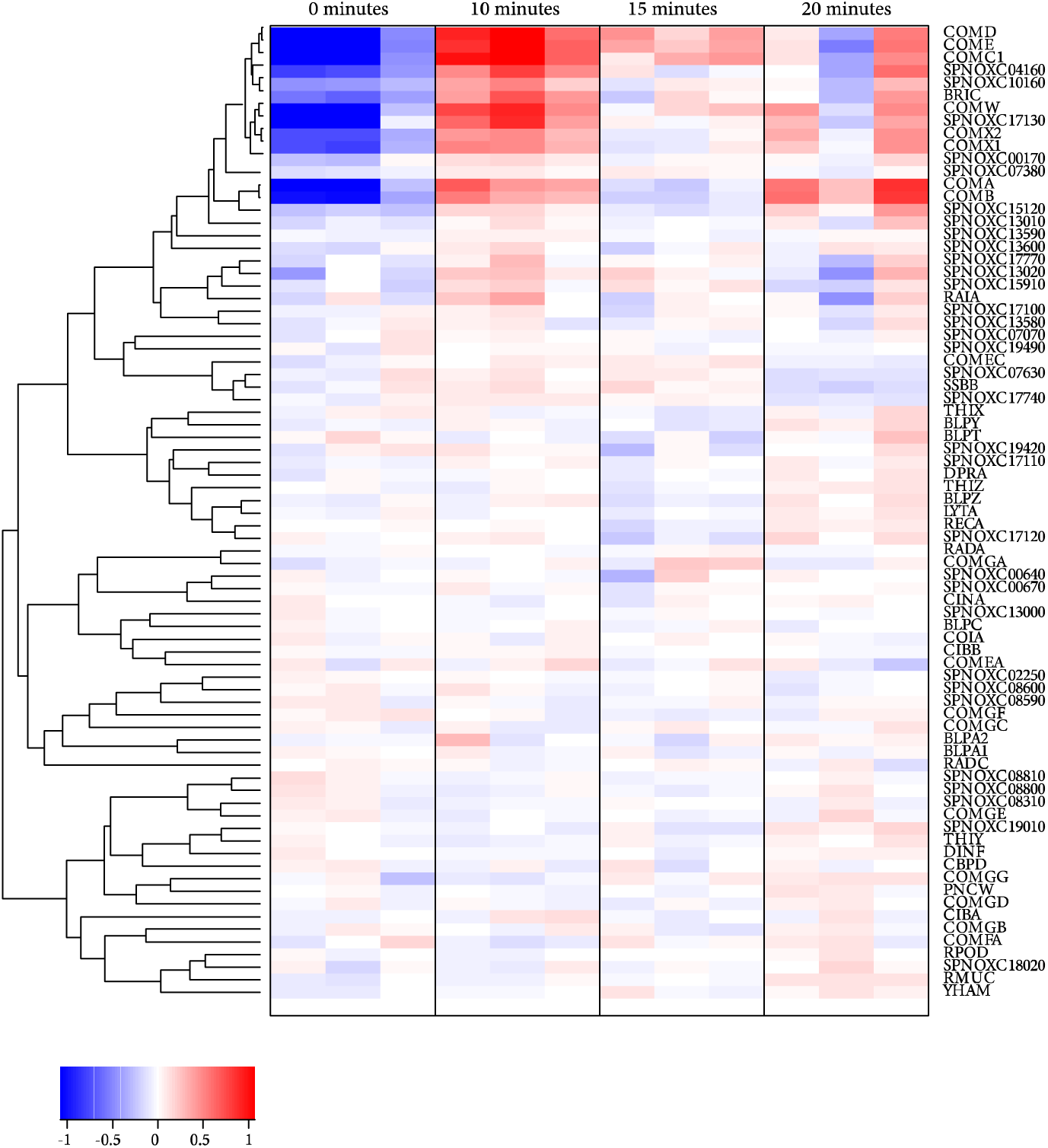
Heatmap showing levels of expression of known competence genes in ND6401 (Clade II) at 10, 15, 20 minutes after CSP addition. Samples are ordered by timepoint and replicate number on the x-axis while genes on y-axis are colored according to their z score and clustered with genes having a similar expression profile using hierarchical clustering. There are three replicates for each timepoint.

Assessing the 15 minute peak-competence timepoint compared to baseline, we observed upregulation of the comAB, comCDE, cibAB and comG operons as well as coding sequences comX and comW. Other notable upregulated genes included SPNOXC16800, a putative single-stranded DNA-binding protein (ssbB); SPNOXC17130, a putative membrane protein proximal to the recA, cinA operon; and SPNOXC1802, a putative methyltransferase located near the competence operon of comYC, a gene essential for genetic transformation. Further, we observed many similarities in expression patterns between the CC180 Clade I strain PT8465 (Clade I) and D39. As expected, we identified upregulation of the comAB, cibAB and comG competence operons as well as genes comC, comD, comX and comW in Clade I strain PT8465. Additionally, we saw consistent upregulation of genes encoding bacteriocins and associated membrane proteins. The bacteriocin peptide briC and co-transcribed membrane protein encoded by SPNOXC04160 are highly upregulated at the 15-minute timepoint with a log2(fold change) of 5.2. Similarly, putative membrane proteins comprising an ABC transporter operon similar to comAB or blpAB operons and encoded by SPNOXC17440, SPNOXC17450, SPNOXC17460 and SPNOXC17470 were found to be upregulated. Notably, SPNOXC17470 is also annotated as a bacteriocin of the Lactococcin 972 family in other S. pneumoniae strains ^35^. CC180 Clade II strain ND6401 demonstrated upregulation of competence genes comAB, comCDE, briC and SPNOXC04160; however, all genes showed a lower level of expression as compared to D39 and PT8465.

**Supplementary Figure 5.**
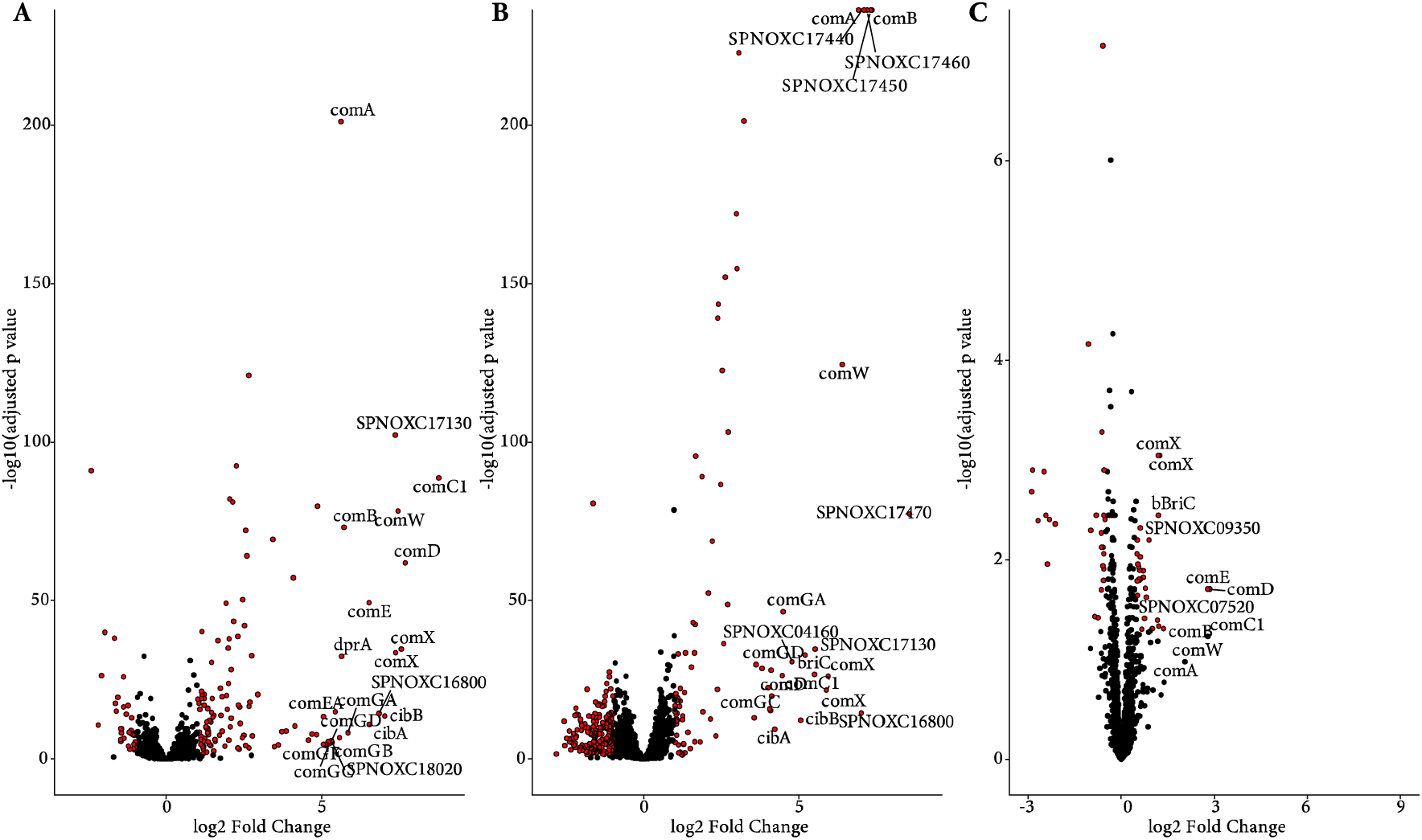
Volcano plot of significantly differentially expressed genes at 15 minutes in strain D39 (panel A), PT8465 (Clade I) (panel B) and ND6401 (Clade II) (panel C). Top 20 genes with highest log2 (fold change) are labelled with gene names. All genes with a p-value less than 0.05 and a log2(fold change) > 1.0 are labelled in red and considered significant while all other genes are shown in black.

### CC180 Clade II strain ND6401 (Clade II) shows a defect in entry into late competence

Through assessment of expression profiles and significant difference in the regulation of competence-associated genes, we identified notable differences in the transition from early to late competence pathways among strains. To further investigate these differences, we plotted the normalized counts and log2(fold change) at 15 minutes compared to baseline for all known essential competence genes. Genes were ordered by timing in the competence pathway beginning with comAB, comCDE operons as well as comX and comW, which are required to initiate entry into the competence state (Supplementary Figures 5A-C). We classified early competence genes as those upregulated by comE including bacteriocins encoded by briC, pncW and the blpABC, blpYZ operons. Late competence genes were classified as those upregulated by comX, which were ordered by position in the genome and clustered by operons.

For D39, we observed upregulation of early competence operons comAB and comCDE as well as bacteriocins. In the late competence genes, we observed that genes encoding proteins involved in DNA repair and recombination such as radA, radC and the recA operon were not highly upregulated. Genes encoding the DNA uptake apparatus comEC, comFA and the comG operon showed consistent high levels of expression at 15 minutes. For the CC180 Clade I strain PT8465, we observed similar transcriptomic changes to D39. Notable differences in the late competence genes included dprA, comFA, comEA and comEC which showed a much smaller log2(fold change) in PT8465. The competence induced protein ccs1 is also downregulated in PT8465 compared to upregulation in D39. We observed the largest differences with our Clade II strain, ND6401. While there was upregulation of the comAB and comCDE operons, comX, comW and briC, we did not observe any upregulation of additional early or late competence genes suggesting a deficit in transitioning to late competence.

**Supplementary Figure 6.**
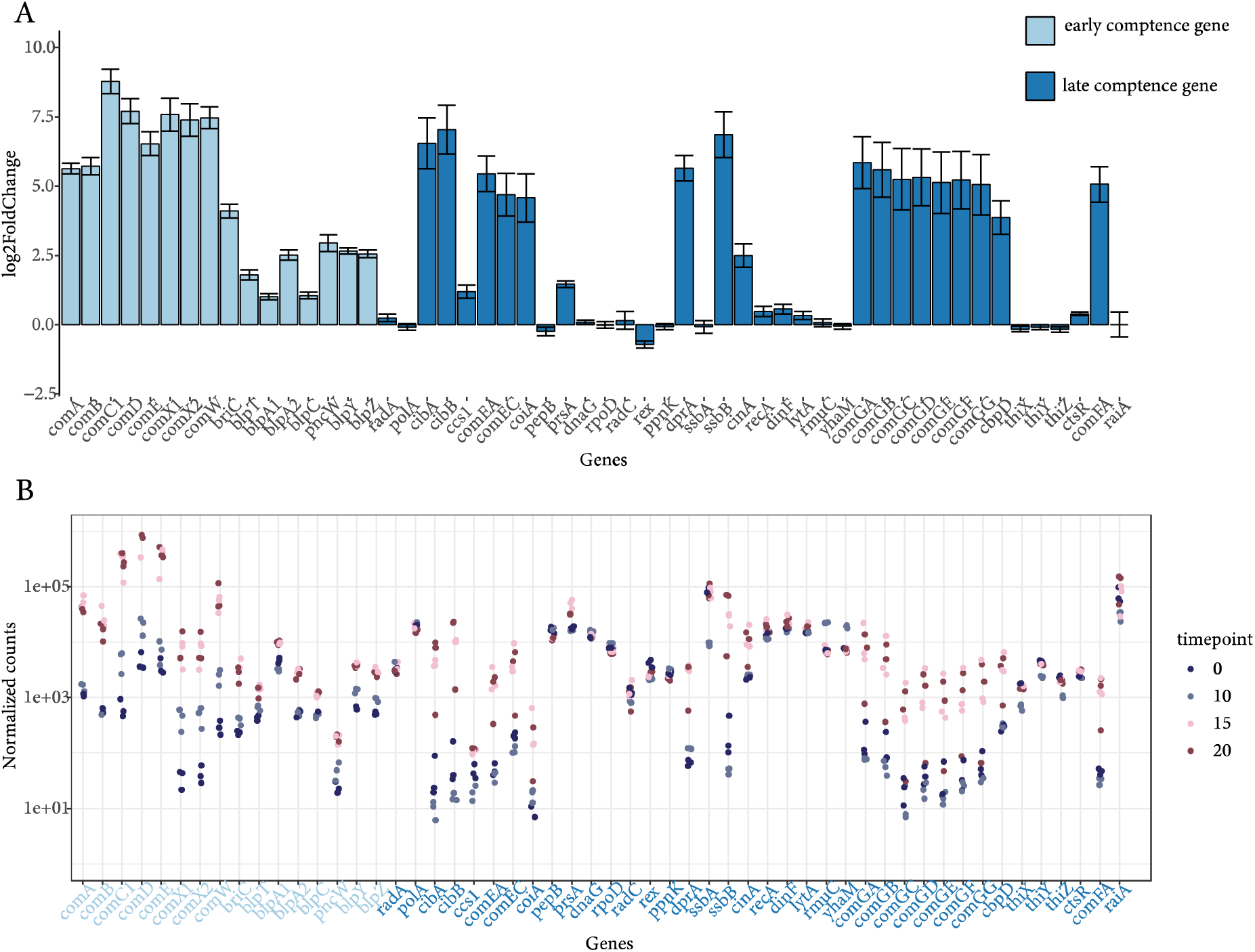
A) Plot of log2(fold change) of competence genes at 15 minutes for D39, as compared to the 0-minute baseline timepoint. Error bars depict standard error of log2(fold change) – lfcSE. All early competence genes are colored in light blue while late competence genes are in dark blue. B) Normalized counts for early and late competence genes showing change in expression levels for D39.

**Supplementary Figure 7.**
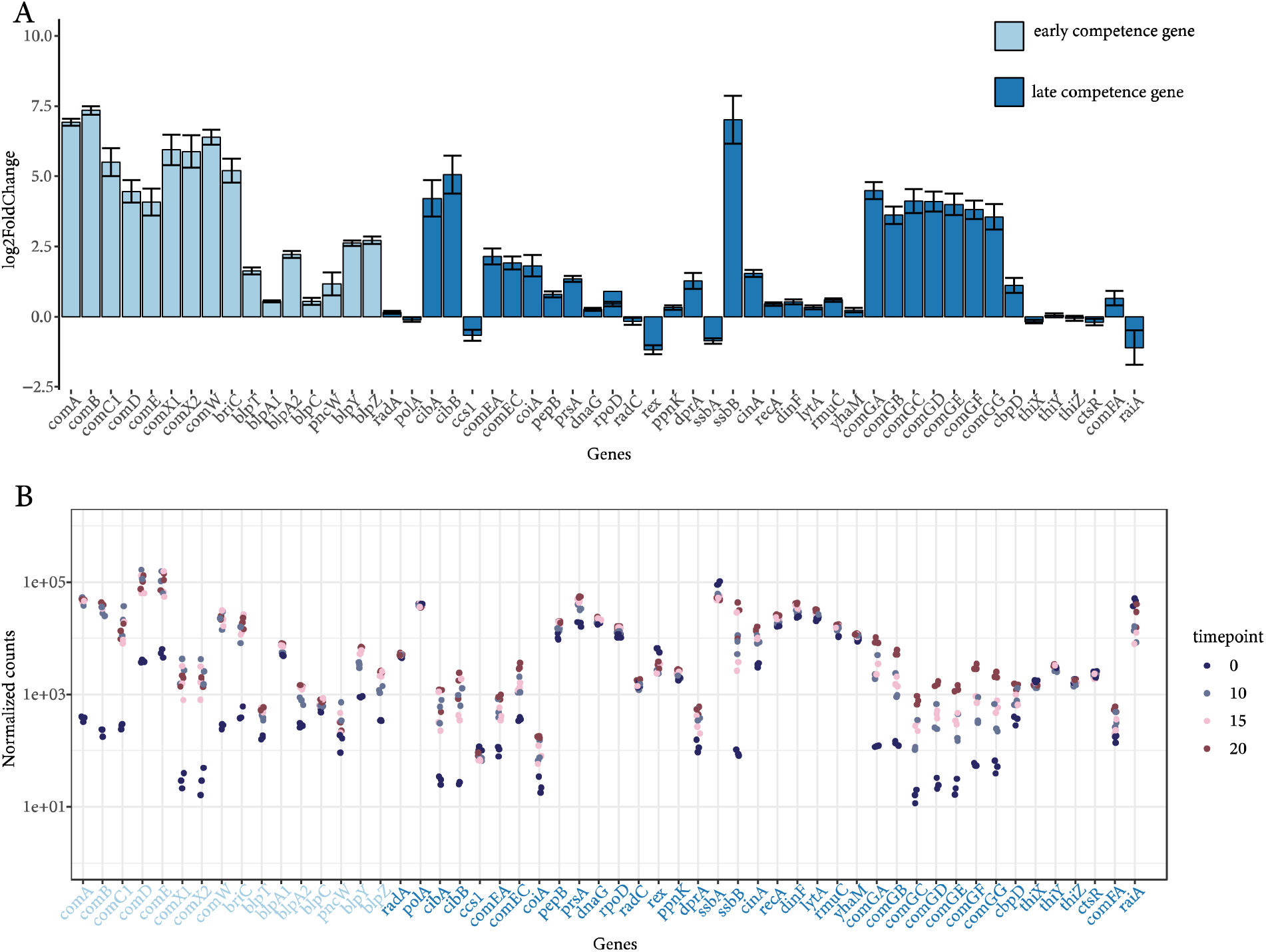
A) Plot of log2(fold change) of competence genes at 15 minutes for PT8465, as compared to the 0-minute baseline timepoint. Error bars depict standard error of log2(fold change) – lfcSE. All early competence genes are colored in light blue while late competence genes are in dark blue. B) Normalized counts for early and late competence genes showing change in expression levels for PT8465.

**Supplementary Figure 8.**
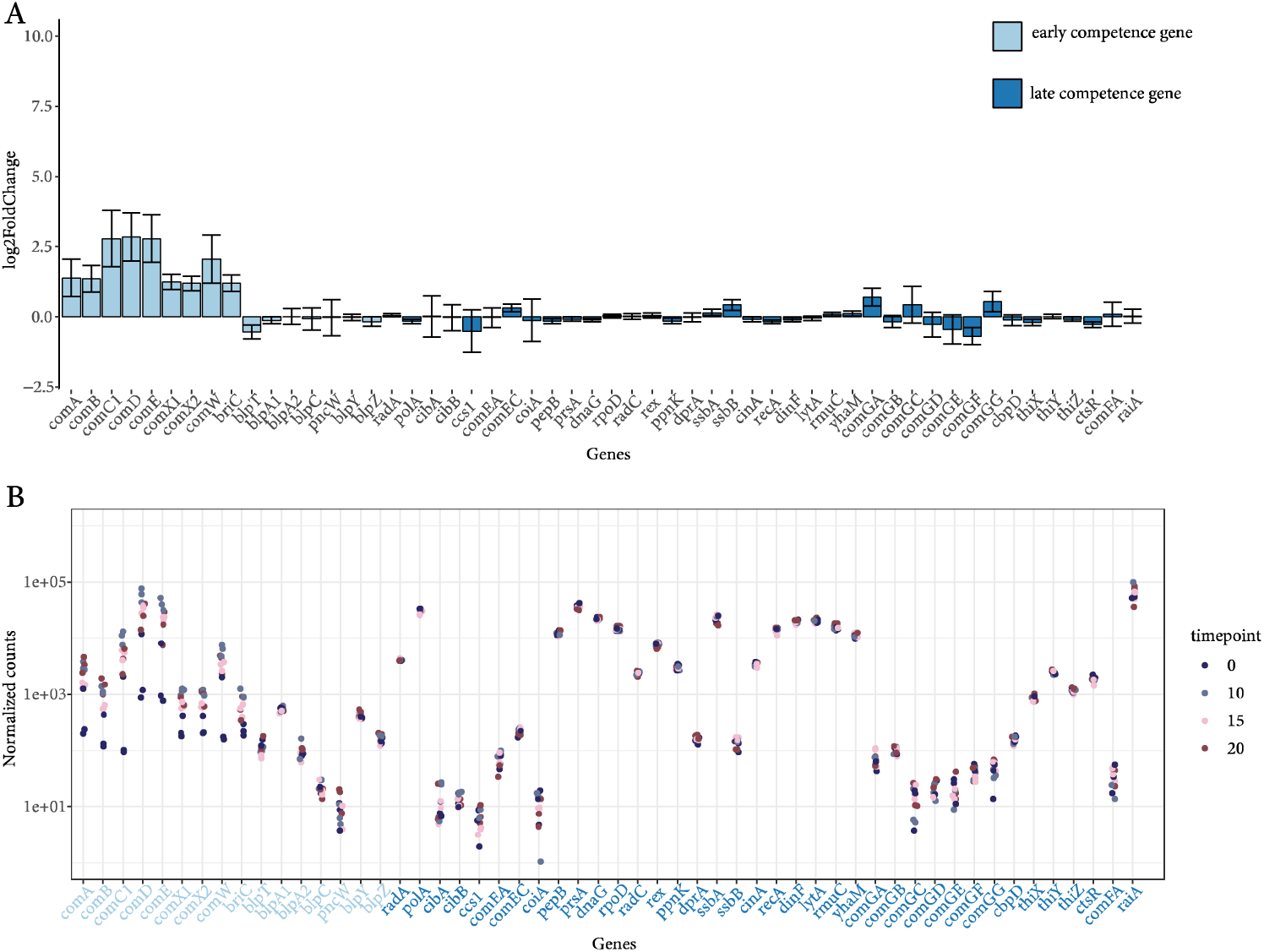
A) Plot of log2(fold change) of competence genes at 15 minutes for ND6401, as compared to the 0-minute baseline timepoint. Error bars depict standard error of log2(fold change) – lfcSE. All early competence genes are colored in light blue while late competence genes are in dark blue. B) Normalized counts for early and late competence genes showing change in expression levels for ND6401.

Last, to further investigate the attenuated expression of genes in the competence pathway seen in Clade II strain ND6401, we compared expression levels of competence genes at baseline (0-minute timepoint prior to CSP addition) for the three strains. We observed a statistically significant higher level of expression of comX1 and comX2 with our Clade II strain ND6401 in comparison to D39 and PT8465 (Supplementary Figure 9). This increased expression prior to competence induction may explain the lack of upregulation seen in late competence genes for this strain which are induced by ComX binding to their promoter regions (Supplementary Figure 9).

**Supplementary Figure 9.**
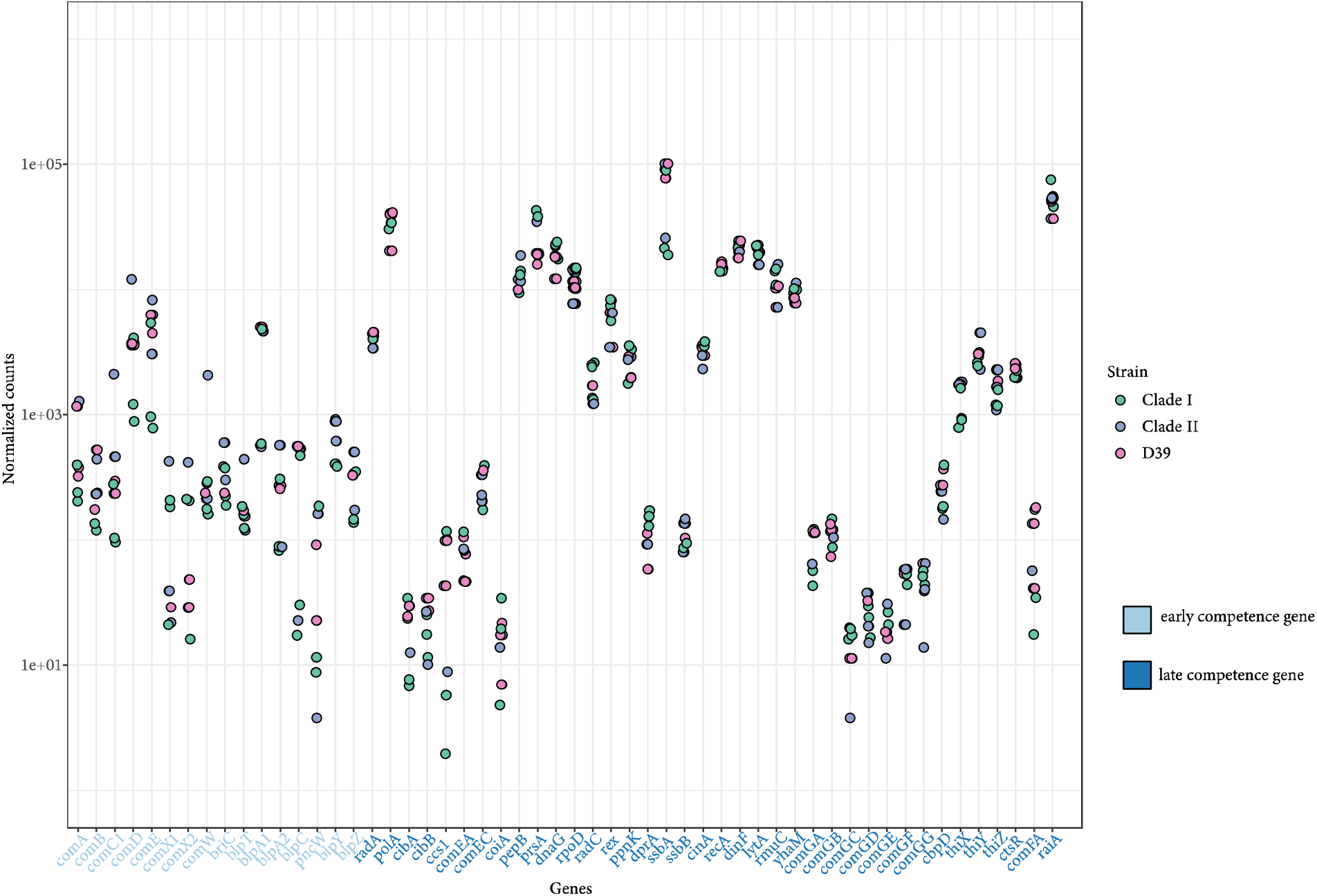
Baseline (0-minute) expression of competence-associated genes among serotype 3 clonal complex 180 strains PT8465 (Clade I) and ND6401 (Clade II), and serotype 2 strain D39. The x-axis lists the genes with early competence genes colored in light blue and late competence genes in dark blue. The y-axis depicts normalized counts of each gene at 0-minutes prior to CSP addition which are colored according to the strain (legend on the right).

## Discussion

S. pneumoniae serotype 3 remains a significant cause of morbidity and mortality, and the recent shift in CC180 population structure has garnered significant interest ^36,37^. Putative contributors to the emergence of Clade II include diversification of non-capsular protein antigens, acquisition of antibiotic resistance, changes in host immunity, or perturbation in pneumococcal population structure associated with vaccination ^10,38^. As several of these factors are directly or indirectly related to competence, understanding variation in recombination rates may elucidate important components of pneumococcal evolution.

Initial genomic investigation of serotype 3 strains belonging to GPSC12/CC180 found that the evolution of Clade I was largely driven by nucleotide substitution ^39^. Further work attributed the reduced recombination rate of pre-PCV13 Clade I strains to the insertion of a 34 Kbp prophage, IZOXC141, which modified a non-coding RNA involved in the induction of transformation ^38^. Analysis of an expanded global sample identified that the emergent Clade II demonstrated a higher rate of recombination, measured by the number of recombination events to polymorphisms introduced by mutation (r/m) using population genomic data. Here, using in-vitro transformation frequency experiments, we found that there is up to two orders of magnitude difference within CC180, with strains belonging to Clade I consistently transforming at a higher rate. This finding contrasts the relative recombination rates of Clades I and II inferred from population genomic analysis. Subsequent transcriptomic analysis revealed that the observed differences in transformation frequency may be due to an attenuated response to CSP among Clade II strains and overall deficiency in the ability to enter the late competence pathway.

Observed rates of bacterial recombination from population genomic samples are the result of both biological and evolutionary processes, mirroring the relationship between mutation rate and molecular evolutionary rate ^40^. Biologically, the bacterium must be naturally competent, possessing the machinery to uptake and incorporate exogenous DNA. Efficiency in this process depends on complex regulation of associated pathways including quorum sensing and bacteriocin production. Additionally, differences in capsular structure, which are associated with strain-specific variation in epidemiology, may impact recombination rates ^39,41^. Larger capsules may physically impede the transfer of DNA across the cell membrane, as unencapsulated pneumococci have higher in-vitro rates of recombination ^42^. Indeed, the mucoid serotype 3 capsule has been shown to contribute to the low recombination rate of the serotype ^38^. Capsule size and charge have also been associated with carriage duration and prevalence, with larger, more negatively charged capsules associated with longer carriage episodes and a higher prevalence due to increased survival from non-opsonic killing by neutrophils ^43,44^. As a result, pneumococci with increased carriage duration may more frequently encounter other streptococci co-colonizing the nasopharynx, leading to more opportunities for homologous recombination^34,45^. Interestingly, serotype 3 is a consistent outlier. Despite it metabolically efficient capsule, extensive capsule production, and negative surface charge, it remains a relatively rare serotype with a shorter carriage duration compared to other serotypes. Recently, new findings suggest that Clade II strains have lower invasiveness, longer carriage duration, and higher bacterial densities in a mouse model compared to Clade I ^46^. If this finding corresponds with human carriage dynamics, then it may partly explain the discordance we observe here between the in-vitro and population genomic measures of recombination. Intriguingly, this suggests that these differences are independent of the capsule.

Entrance into competence corresponds with a shift in the transcriptional profile of the bacterial cell, beginning with the signal from CSP. Previous transcriptomic analysis of serotype 3 CC180 strain OXC141 found that the competence-inducing comCDE operon, which encodes the CSP precursor and its receptor, is transcribed in an antisense manner ^39^. Indeed, all serotype 3 strains studied here had antisense transcription of the operon. In addition, among Clade I strains harboring φOXC141, the prophage was found to modify csRNA3, a non-coding RNA that inhibits the induction of transformation ^38^. However, while this explain in part the low in-vivo recombination of φOXC141-postive Clade I strains, a direct comparison to Clade II strains was not performed. In the present study, only one of the Clade I strains tested (biosample SAMN37512611) harbored the IZOXC141 prophage, and this strain showed comparable transformation frequencies to IZOXC141 negative Clade I strains, albeit with a larger confidence interval. Nevertheless, isolates from both clades were able to undergo transformation. We do observe that expression of late competence genes in the representative Clade II strain was attenuated, suggesting defects in transitioning to stages of late competence. However, further detailed investigation of promoter sequences for related operons is needed to explain fine-tuned transcriptional control of competence pathways among serotype 3.

Other notable differences include the upregulation of multiple bacteriocins among serotype 3 strains but not D39. Bacteriocins, toxic peptides involved in bacterial fratricide, are a diverse group of proteins which mediate inter-strain competition ^47^. Differences in expression of bacteriocin genes may explain the population level differences seem among our strains ^48^. Strains with higher expression of bacteriocins are believed to fare better against selective pressure in the environment. Previous work has also shown that expression of S. pneumoniae bacteriocins is induced by antibiotics via regulatory interplay with the competence system ^49^. This may also explain differences in transformation frequency between closely related strains due to genomic differences in bacteriocin genes. Taken together, differences in the composition and expression of bacteriocins among strains belonging to Clades I and II may dually explain both the longer carriage duration and discordance in in-vitro and population genomic rates of recombination. In other words, while Clade II strains have lower in-vitro transformation efficiency, increased carriage duration and upregulation of bacteriocins may have resulted in increased in-vivo recombination success, as observed in population genomic data. This warrants further investigation.

We acknowledge some limitations of the in-vitro transformation frequency assay as a measure of recombination rate. Using saturating concentrations of donor DNA in the form of a PCR amplified fragment of the kanamycin resistance gene aphIII, which has a product size of 977 bp, does not allow us to observe differences in tract length of recombinant DNA. In prior studies, the largest single recombination event observed using comparisons of genome sequences ranged from 20,000 – 30,000 bp ^50^. This may be due to the fact that in vivo, streptococci gain more diversity through the recombination of a small number of large size DNA fragments released from the genomic DNA of nearby bacterial cells undergoing fratricide. For further experiments, it would be interesting to use ultra-long read Oxford Nanopore sequencing to map the recombination of large tracks of DNA.

Understanding the mechanisms behind variation in horizontal gene transfer for S. pneumoniae can contribute to understanding the impact recombination has on bacterial species including their ability to diversify and adapt to human interventions. Our observations also indicate that transformation frequency is not solely dictated by serotype and may vary significantly even among closely related strains.

## References

1. Geno KA, Gilbert GL, Song JY, et al. Pneumococcal Capsules and Their Types: Past, Present, and Future. Clin Microbiol Rev. 2015;28(3):871-899. doi:10.1128/CMR.00024-15

2. Ahl J, Littorin N, Forsgren A, Odenholt I, Resman F, Riesbeck K. High incidence of septic shock caused by Streptococcus pneumoniae serotype 3 - a retrospective epidemiological study. BMC Infect Dis. 2013;13(1):492. doi:10.1186/1471-2334-13-492

3. Martens P, Worm SW, Lundgren B, Konradsen HB, Benfield T. Serotype-specific mortality from invasive Streptococcus pneumoniaedisease revisited. BMC Infect Dis. 2004;4(1):21. doi:10.1186/1471-2334-4-21

4. Ganaie F, Saad JS, McGee L, et al. A new pneumococcal capsule type, 10D, is the 100th serotype and has a large cps fragment from an oral streptococcus. mBio. 2020;11(3). doi:10.1128/MBIO.00937-20/SUPPL_FILE/MBIO.00937-20-SF005.TIF

5. Subramanian R, Liyanapathirana V, Barua N, et al. Persistence of pneumococcal serotype 3 in adult pneumococcal disease in hong kong. Vaccines (Basel). 2021;9(7). doi:10.3390/vaccines9070756

6. Horácio AN, Silva-Costa C, Lopes E, Ramirez M, Melo-Cristino J. Conjugate vaccine serotypes persist as major causes of non-invasive pneumococcal pneumonia in Portugal despite declines in serotypes 3 and 19A (2012-2015). PLoS One. 2018;13(11). doi:10.1371/journal.pone.0206912

7. Forstner C, Kolditz M, Kesselmeier M, et al. Pneumococcal conjugate serotype distribution and predominating role of serotype 3 in German adults with community-acquired pneumonia. Vaccine. 2020;38(5):1129–1136. doi:10.1016/j.vaccine.2019.11.026

8. Cella E, Sutcliffe CG, Grant LR, et al. Streptococcus pneumoniae serotype 3 population structure in the era of conjugate vaccines, 2001–2018. Microb Genom. 2024;10(3):001196. doi:10.1099/MGEN.0.001196/CITE/REFWORKS

9. Groves N, Sheppard CL, Litt D, et al. Evolution of Streptococcus pneumoniae Serotype 3 in England and Wales: A Major Vaccine Evader. Genes 2019, Vol 10, Page 845. 2019;10(11):845. doi:10.3390/GENES10110845

10. Azarian T, Mitchell PK, Georgieva M, et al. Global emergence and population dynamics of divergent serotype 3 CC180 pneumococci. Mitchell TJ, ed. PLoS Pathog. 2018;14(11):e1007438. doi:10.1371/journal.ppat.1007438

11. Croucher NJ, Harris SR, Fraser C, et al. Rapid Pneumococcal Evolution in Response to Clinical Interventions. Science (1979). 2011;331. http://science.sciencemag.org/

12. Golubchik T, Brueggemann AB, Street T, et al. Pneumococcal genome sequencing tracks a vaccine escape variant formed through a multi-fragment recombination event. Nat Genet. 2012;44(3):352. doi:10.1038/NG.1072

13. Mostowy RJ, Croucher NJ, De Maio N, et al. Pneumococcal Capsule Synthesis Locus cps as Evolutionary Hotspot with Potential to Generate Novel Serotypes by Recombination. Mol Biol Evol. 2017;34(10):2537–2554. doi:10.1093/molbev/msx173

14. Croucher NJ, Kagedan L, Thompson CM, et al. Selective and genetic constraints on pneumococcal serotype switching. PLoS Genet. 2015;11(3):e1005095. doi:10.1371/journal.pgen.1005095

15. Lin J, Zhu L, Lau GW. Disentangling competence for genetic transformation and virulence in Streptococcus pneumoniae. Curr Genet. 2016;62(1):97–103. doi:10.1007/s00294-015-0520-z

16. Kilian M, Poulsen K, Blomqvist T, et al. Evolution of Streptococcus pneumoniae and its close commensal relatives. PLoS One. 2008;3(7). doi:10.1371/journal.pone.0002683

17. Chaguza C, Cornick JE, Everett DB. Mechanisms and impact of genetic recombination in the evolution of Streptococcus pneumoniae. Comput Struct Biotechnol J. 2015;13:241–247. doi:10.1016/j.csbj.2015.03.007

18. Mostowy R, Croucher NJ, Hanage WP, Harris SR, Bentley S, Fraser C. Heterogeneity in the frequency and characteristics of homologous recombination in pneumococcal evolution. PLoS Genet. 2014;10(5):e1004300. doi:10.1371/journal.pgen.1004300

19. Joloba ML, Kidenya BR, Kateete DP, et al. Comparison of transformation frequencies among selected Streptococcus pneumoniae serotypes. Int J Antimicrob Agents. 2010;36(2):124–128. doi:10.1016/j.ijantimicag.2010.03.024

20. Evans BA, Rozen DE. Significant variation in transformation frequency in Streptococcus pneumoniae. ISME Journal. 2013;7(4):791–799. doi:10.1038/ismej.2012.170

21. Hsieh YC, Wang JT, Lee W sen, et al. Serotype competence and penicillin resistance in Streptococcus pneumoniae. Emerg Infect Dis. 2006;12(11):1709–1714. doi:10.3201/eid1211.060414

22. Pestova E V., Håvarstein LS, Morrison DA. Regulation of competence for genetic transformation in Streptococcus pneumoniae by an auto-induced peptide pheromone and a two-component regulatory system. Mol Microbiol. 1996;21(4):853–862. doi:10.1046/J.1365-2958.1996.501417.X

23. Cheng Q, Campbell EA, Naughton AM, Johnson S, Masure HR. The com locus controls genetic transformation in Streptococcus pneumoniae. Mol Microbiol. 1997;23(4):683–692. doi:10.1046/J.1365-2958.1997.2481617.X

24. Ween O, Gaustad P, Sigve L, Varstein HÊ. Identification of DNA binding sites for ComE, a key regulator of natural competence in Streptococcus pneumoniae. Mol Microbiol. 1999;33(4):817–827.

25. Luo P, Morrison DA. Transient association of an alternative sigma factor, ComX, with RNA polymerase during the period of competence for genetic transformation in Streptococcus pneumoniae. In: Journal of Bacteriology. Vol 185. ; 2003:349–358. doi:10.1128/JB.185.1.349-358.2003

26. Martin B, Garcfa P, Castanie MP, Claverys’ • JP. The recA gene of Streptococcus pneumoniae is part of a competence-induced operon and controls lysogenic. Mol Microbiol. 1995;15(2):367–379.

27. Claverys JP, Martin B, Håvarstein LS. Competence-induced fratricide in streptococci. Mol Microbiol. 2007;64(6):1423–1433. doi:10.1111/j.1365-2958.2007.05757.x

28. Weng L, Piotrowski A, Morrison DA. Exit from Competence for Genetic Transformation in Streptococcus pneumoniae Is Regulated at Multiple Levels. PLoS One. 2013;8(5). doi:10.1371/journal.pone.0064197

29. Martin B, Soulet AL, Mirouze N, et al. ComE/ComE∼P interplay dictates activation or extinction status of pneumococcal X-state (competence). Mol Microbiol. 2013;87(2):394–411. doi:10.1111/mmi.12104

30. Peterson SN, Sung CK, Cline R, et al. Identification of competence pheromone responsive genes in Streptococcus pneumoniae by use of DNA microarrays. Mol Microbiol. 2004;51(4):1051–1070. doi:10.1046/J.1365-2958.2003.03907.X

31. Peterson S, Cline RT, Tettelin H, Sharov V, Morrison DA. Gene expression analysis of the Streptococcus pneumoniae competence regulons by use of DNA microarrays. J Bacteriol. 2000;182(21):6192–6202. doi:10.1128/JB.182.21.6192-6202.2000

32. Slager J. Triggering pneumococcal competence: Memoirs of an escape artist. Published online 2019. Accessed July 7, 2024. https://research.rug.nl/en/publications/triggering-pneumococcal-competence-memoirs-of-an-escape-artist

33. Shambhu S, Cella E, Jubair M, Azarian T. Complete Genome Sequences of Nine Streptococcus pneumoniae Serotype 3 Clonal Complex 180 Strains. Microbiol Resour Announc. 2022;11(7). doi:10.1128/MRA.00275-22

34. Li G, Liang Z, Wang X, et al. Addiction of Hypertransformable Pneumococcal Isolates to Natural Transformation for In Vivo Fitness and Virulence. Infect Immun. 2016;84(6):1887–1901. doi:10.1128/IAI.00097-16

35. Martínez B, Suárez JE, Rodríguez A. Lactococcin 9721: a homodimeric lactococcal bacteriocin whose primary target is not the plasma membrane. Microbiology (Reading). 1996;142 (Pt 9)(9):2393-2398. doi:10.1099/00221287-142-9-2393

36. Luck JN, Tettelin H, Orihuela CJ. Sugar-Coated Killer: Serotype 3 Pneumococcal Disease. Front Cell Infect Microbiol. 2020;10(December):1–11. doi:10.3389/fcimb.2020.613287

37. Sings HL, Gessner BD, Wasserman MD, Jodar L. Pneumococcal Conjugate Vaccine Impact on Serotype 3: A Review of Surveillance Data. Infect Dis Ther. 2021;10(1):521–539. doi:10.1007/s40121-021-00406-w

38. Kwun MJ, Ion A V., Cheng HC, et al. Post-vaccine epidemiology of serotype 3 pneumococci identifies transformation inhibition through prophage-driven alteration of a non-coding RNA. Genome Med. 2022;14(1):1–23. doi:10.1186/S13073-022-01147-2/FIGURES/4

39. Croucher NJ, Mitchell AM, Gould KA, et al. Dominant role of nucleotide substitution in the diversification of serotype 3 pneumococci over decades and during a single infection. PLoS Genet. 2013;9(10):e1003868. doi:10.1371/journal.pgen.1003868

40. Croucher NJ, Coupland PG, Stevenson AE, Callendrello A, Bentley SD, Hanage WP. Diversification of bacterial genome content through distinct mechanisms over different timescales. Nat Commun. 2014;5(March 2016):1–12. doi:10.1038/ncomms6471

41. Croucher NJ, Finkelstein JA, Pelton SI, et al. Population genomics of post-vaccine changes in pneumococcal epidemiology. Nat Genet. 2013;45(6):656–663. doi:10.1038/ng.2625

42. Yother J, McDaniel LS, Briles DE. Transformation of encapsulated Streptococcus pneumoniae. J Bacteriol. 1986;168(3):1463–1465. doi:10.1128/JB.168.3.1463-1465.1986

43. Weinberger DM, Trzcinski K, Lu YJ, et al. Pneumococcal capsular polysaccharide structure predicts serotype prevalence. Bishai W, ed. PLoS Pathog. 2009;5(6):e1000476. doi:10.1371/journal.ppat.1000476

44. Li Y, Weinberger DM, Thompson CM, Trzciński K, Lipsitch M. Surface charge of Streptococcus pneumoniae predicts serotype distribution. Infect Immun. 2013;81(12):4519–4524. doi:10.1128/IAI.00724-13

45. Chaguza C, Andam CP, Harris SR, et al. Recombination in Streptococcus pneumoniae lineages increase with carriage duration and size of the polysaccharide capsule. mBio. 2016;7(5):e01053–16. doi:10.1128/mBio.01053-16

46. Sekulovic O, Gallagher C, Lee J, et al. Evidence of Reduced Virulence and Increased Colonization among Pneumococcal Isolates of Serotype 3 Clade II Lineage in Mice. J Infect Dis. Published online January 29, 2024. doi:10.1093/INFDIS/JIAE038

47. Dawid S, Roche AM, Weiser JN. The blp bacteriocins of Streptococcus pneumoniae mediate intraspecies competition both in vitro and in vivo. Infect Immun. 2007;75(1):443–451. doi:10.1128/IAI.01775-05

48. Lehtinen S, Croucher NJ, Blanquart F, Fraser C. Epidemiological dynamics of bacteriocin competition and antibiotic resistance. Proceedings of the Royal Society B. 2022;289(1984). doi:10.1098/RSPB.2022.1197

49. Kjos M, Miller E, Slager J, et al. Expression of Streptococcus pneumoniae Bacteriocins Is Induced by Antibiotics via Regulatory Interplay with the Competence System. PLoS Pathog. 2016;12(2):e1005422. doi:10.1371/JOURNAL.PPAT.1005422

50. Cowley LA, Petersen FC, Junges R, Jimson D. Jimenez M, Morrison DA, Hanage WP. Evolution via recombination: Cell-to-cell contact facilitates larger recombination events in Streptococcus pneumoniae. PLoS Genet. 2018;14(6):e1007410. doi:10.1371/JOURNAL.PGEN.1007410

